# *Veillonella parvula*: a strictly anaerobic bacterium with high efficacy for safe and specific tumor targeting and colonization

**DOI:** 10.1101/2021.05.10.443531

**Authors:** Amirhosein Kefayat, Fatemeh Ghahremani, Soodabeh Rostami

**Author notes:** **Corresponding author:** *Fatemeh Ghahremani*, Postal code: 38481-76941, Address: Sardasht, Meydan Basij, Arak University of Medical Sciences.

## Abstract

Bacterial cancer therapy has gained lots of attention in the past decade and is now considering a reliable option for the future. However, some concerns have limited its application into clinic settings like insufficient colonization of tumors and infectious origin of the currently used bacteria like Clostridium and Salmonella species, especially in cancer patients which exhibit different levels of immunocompromising. In the present study, *Veillonella parvula* (*V. parvula*) as a strictly anaerobic bacterium which has rarely identified as a pathogen in human, was administrated into 4T1 breast tumor-bearing mice. At first, 4T1 breast tumor-bearing BALB/c mice were injected with 10^7^ bacteria intravenously, intraperitoneally, orally, or intratumorally. The best administration route according to tumor colonization and safety was selected. Then, the therapeutic effect of *V. parvula* administration was investigated according to the 4T1 breast tumor’s growth, metastasis, and tumor-bearing mice survival. Besides, histopathological evaluations were done to estimate microscopic changes at the inner of the tumor. *V. parvula* exhibited significant tumor-targeting and colonization efficacy, 24 h after intravenous administration and formed clustered colonies at the central region of the tumors. Although a negligible number of the bacteria were localized at normal organs, these organs became clear from the bacteria after 72 h, and no side effects or death were observed at the animals after intravenous administration of *V. parvula*. Although mean tumor volumes in the *V. parvula* treated group was lower than the control (~ 25.4%), their difference wasn’t statistically significant (*P* > 0.05). Despite significant tumor colonization (5500000:1 in comparison with normal organs after 72 h), *V. parvula* didn’t cause a significant therapeutic effect on the metastasis or survival time of tumor-bearing mice. Taking together, *V. parvula* is a completely safe and tumor-specific agent per se, without any genetic manipulation. Also, it exhibits high tumor penetration and colonization at the deep regions of the tumor.

## 1. Introduction

Bacteria as the old enemies of human societies were the main causes of human death in the past. However, their age of power declined by the development of antibiotics, vaccination, and enhancement of hygiene^1^. Nowadays cancers are one of the main causes of death, worldwide. Some observations exhibited that bacteria can cause significant therapeutic effects on the tumor in different aspects^2^. William B. Coley was the pioneer in employing bacteria for cancer therapy. He utilized alive *Streptococcus pyogenes* for treatment of a head and neck cancer patient which caused hopeful results^3^. Subsequently, over one thousand cancer patients were treated with Coley’s toxins and excellent results were observed^4,5^. The rise of radiation therapy at that time caused the fading of the bacteria therapy outcomes. But, the resistance of cancer cells to the current treatments and their extensive side effects made researchers focus on the new or even forgotten therapeutic approaches like bacteria therapy for cancer treatment^6–8^.

The tumor is a hospitable microenvironment for bacteria localization and colonization. Bacteria can hide from the immune system at the tumor site^9^ and the tumor microenvironment can provide enough nutrition for them^10^. Tumor hypoxia is one of the main predisposing factors for tumor colonization of bacteria^11,12^. Therefore, anaerobic species like *Escherichia*^13^, *Clostridium*^14^, *Bifidobacterium*^15,16^, *Salmonella*^17^, and *Listeria*^18^ have gained lots of attention for cancer treatment due to tumor-specific colonization after systemic administration. Also, tumor hypoxic regions are the main site of chemo and radiation therapy resistance. These regions have a determinative effect on the patients’ prognosis^19^. Bacteria colonization at the tumor site can cause anti-tumor effects through different mechanisms including activating the immune system and recruitment of immune cells to the tumor^20,21^, production of anti-tumor substances^6^, induction of cancer cells apoptosis due to intracellular proliferation, and competition for nutrition resources^22^.

Many different bacteria species have exhibited a high ability for colonizing solid tumors which surprisingly caused anti-tumor effects. Some of the used species are pathogenic which can kill the host if left untreated. Therefore, their attenuated subtypes were designed by genetic manipulation for *in vivo* experiments^4,23^.

In the present study, we hypothesized to use a strictly anaerobic bacteria which is highly vulnerable against oxygen^24^ for tumor targeting and colonization. Facultative anaerobic bacteria like *Salmonella* are not limited to hypoxia and can colonize normal tissues which is a negative point for utilizing this species for tumor treatment^25^. However, strict anaerobic property can limit the administered bacteria localization and colonization just at the tumor site and specially at the hypoxic regions of the tumor which are mostly out of access for other treatments^11^. Also, *V. parvula* is not pathogenic, and extremely rare case studies have reported the *V. parvula* caused infections^26,27^. *V. parvula* uptakes lactate as a determinative energy resource for cancer cells and mainly produces propionate^28^ which has anti-tumor effects^29^. Therefore, its therapeutic effects on tumor growth and metastasis was evaluated in the 4T1 murine breast cancer model in BALB/c mice.

## 2. Method & materials

### 2.1. Bacterium culture

*V. parvula* (ATCC 10790) was cultured on Reinforced Clostridial medium (Oxoid CM149) with sodium lactate (60% solution) at a concentration of 1.5% at 37 °C in the anaerobic atmosphere (gas mixture, 80% N2-10% CO2-10% H2) for 5 days.

### 2.2. Cancer cells culture

Mouse mammary carcinoma cell line (4T1) was purchased from Pasteur Institute of Tehran, Iran. Cells were cultured in RPMI 1640 medium (Sigma, USA) containing 10% fetal bovine serum (FBS) (Sigma, USA) and 1% antibiotics mixture containing penicillin (Sigma, USA) and streptomycin (Sigma, USA). The cells were incubated at 37 ° C in a humidified incubator at a 5% CO_2_ atmosphere.

### 2.3. Animals husbandry and handling

Female BALB/c mice (mean weight: 25 ± 2 gr) were purchased from the Pasteur Institute of Tehran, Iran. The mice were maintained at 24 ± 2 °C temperature, 50 ±10% relative humidity, and 12 h light/12 h dark cycle condition with complete access to standard mouse chow and water. The mice were acclimated for at least 2 weeks before the start of the study. If any signs of pain, wounds, massive necrosis and hemorrhage, diffuse metastasis were observed during any steps of the study, the mice were sacrificed by Ketamine/Xylazine solution overdose.

### 2.4. 4T1 cancer cell implantation and tumor-bearing mice monitoring

BALB/c mice were injected subcutaneously (s.c) into the 4^th^ abdominal mammary fat pad with 1 × 10^6^ 4T1 cells suspended in 50 μl PBS. The injection site was shaved and sterilized with 70% alcohol before injection. To determine tumors’ growth progression, the greatest longitudinal diameter (length) and the greatest transverse diameter (width) of the tumors were determined every 3 days for 27 days after the cancer cells implantation. Then, the tumor’s volume was calculated by the tumor volume equation (1)^30^.

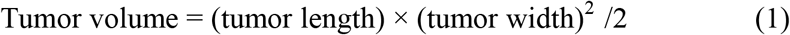

For survival analyzes, tumor-bearing mice survival were monitored for 60 days after the cancer cells implantation and Kaplan-Meier analyzes was used to compare the difference between the groups’ survival time. If any signs of pain, wounds, massive tumor necrosis, hemorrhage, or diffuse metastasis were observed during any steps of the study, the mice were sacrificed by an overdose of ketamine/xylazine solution. It should be mentioned that standardized humane endpoints based on the current guidelines for endpoints in animal tumor studies were used^31–33^.

### 2.5. Bacteria biodistribution

To evaluate the bacteria biodistribution in tumor-bearing mice body through different routes of administration, 10^7^ *V*. parvula were intraperitoneally (i.p), intravenously (i.v), intratumorally (i.t), and orally (p.o) administered into the tumor-bearing mice (tumor volume: 100 ± 34 mm^3^). Each group contained 10 mice (n=10). 5 mice were sacrificed after 48 and the remaining 5 were sacrificed after 72 h from the administration. The tissue samples from the tumor, liver, lungs, spleen, and kidneys were harvested, weighed, and then 0.1 g of each tissue was homogenized in 3 ml of PBS at room temperature for 1 minute in a completely aseptic condition to extract the bacteria from tissues. Then, to determine the presence of the bacteria, 10 μl homogenized tissues were plated onto the standard culture medium for *V. parvula* and incubated 72 h in anaerobic condition and bacterial colony count was evaluated in colony-forming units (CFUs).

### 2.6. Histopathology exams

The harvested organs were fixed in 10% formalin neutral buffer solution for at least 24 h. The fixed specimens were processed overnight for dehydration, clearing, and impregnation using an automatic tissue processor (Sakura, Japan). The specimens were embedded in paraffin blocks and serial sections of 4 μm thickness were cut using a microtome (Leica Biosystems, Germany). The sections were stained by Hematoxylin & Eosin (H&E)^34^ Histological photographs were obtained using a digital light microscope (Olympus, Japan).

### 2.6. Blood biochemical analyzes

16 healthy female BALB/c mice were randomly divided into 2 groups (n=5) including PBS and *V. parvula* groups. The mice in the 1^st^ group were injected with PBS. The 2^nd^ group was i.v injected with 10^7^ bacteria. The mice were closely monitored for mortality, appearance, behavioral pattern changes such as weakness, aggressiveness, food or water refusal, and pain or any signs of illness within these 30 days. On the 30^th^ day, the mice were sacrificed and their blood was collected and the discarded serum was used for biochemistry evaluations. Creatinine (Cr), alanine aminotransferase (ALT), and aspartate aminotransferase (AST) were measured for biochemical assays. Moreover, their organs were harvested and fixed in 10% formalin neutral buffer solution for histopathological analyzes.

### 2.8. Statistical analyses

The statistical analyses were performed using one-way analysis of variance (ANOVA) with Tukey’s post-hoc test by JMP 11.0 software (SAS Institute, Japan). Statistical significance was set at P < 0.05 (*: P < 0.05, ns: not significant). The results were illustrated as mean ± standard deviation (SD). All experiments were replicated at least three times.

### 2.9. Ethics statement

All animal experiments complied with the ARRIVE guidelines and were carried out in accordance with the guidelines of the European Communities Council Directive (2010/63/UE). Also, all the procedures were approved by the Ethics Committee for Animal Experimentation of the Arak University of Medical Sciences (IR.ARAKMU.REC.1398.012).

## 3. Results & Discussion

### 3.1. Tumor targeting efficacy of the *V. parvula* after systemic administration

One of the most determinative factors for the efficacy of a bacterium species to be used for tumor targeting and colonization is its tumor specificity^35^. It’s obvious that more tumor colonization and less preference for the normal organs localization can significantly enhance the tumor-targeting efficacy and decrease the side effects^36^. The administration route can significantly affect tumor colonization efficacy. Four common routes including intraperitoneal (i.p) injection, intravenous injection (i.v), intratumoral (i.t) injection and oral gavage (p.o) have been utilized for the bacteria administration in different studies^37^. Therefore, 1×10^7^ *V. parvula* was administrated i.p, i.v, p.o, and i.t to the 4T1 breast tumor-bearing BALB/c mice to identify the best route of *V. parvula* administration. Tumors had 100 ± 34 mm^3^ volume on the administration day. Tissue samples from the tumor, liver, lungs, spleen, and kidneys of the tumor-bearing mice were collected 48 and 72 h after administration and biodistribution of the bacteria was evaluated (Figure 1). As Figure 1A and B illustrate, i.v administration of *V. parvula* caused the highest efficacy for tumor colonization in comparison with i.p injection, p.o, and even i.t injection (Figure 1A) at both time points. Although i.v administration caused *V. parvula* localization at the normal organs, it was temporary and the bacteria were eradicated from these organs after 72 h (Figure 1B). The lowest efficacy according to tumor colonization was observed at p.o administered tumor-bearing mice. No bacteria growth was detected at the tumor of p.o administered tumor-bearing mice. The other significant weakness of p.o administration was a 60% mortality rate in the next five days after the *V. parvula* administration (Figure 1C) which was due to aspiration and forming lung abscess lesions according to histopathological evolutions (Figure 1D & E). Also. lungs of these dead mice were fulfilled by macrophages (Figure 1F). On the other side, no sign of fever, chills, cachexia, anorexia, and body weight loss or death was observed in other groups during these 30 days of monitoring.

**Figure 1:**
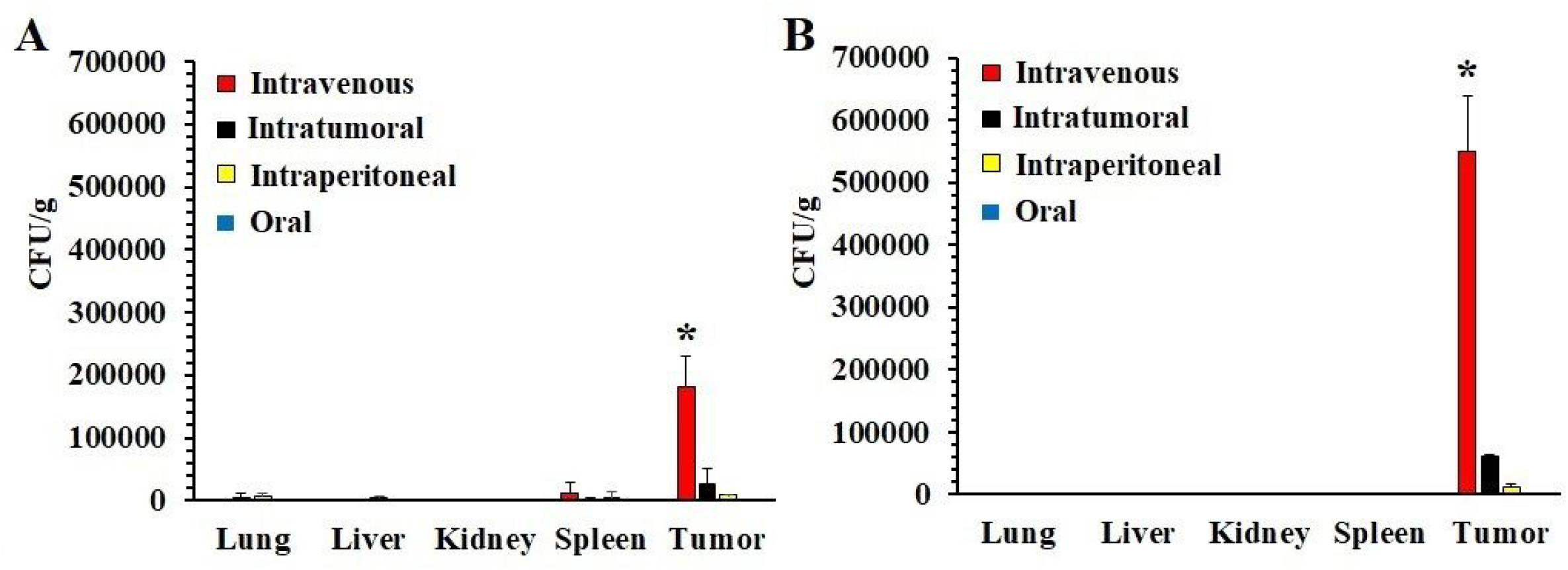
Efficacy of different routes of *V. parvula* administration according to the bacteria biodistribution and tumor colonization at the 4T1 breast tumor-bearing mice (**A**) 48 h (n=5) and (**B**) 72 h (n=5) after 10 *V. parvula* administration. **(C)** Kaplan-Meier survival chart of mice after *V. parvula* administration through different routes (n=8). (**D**) Macroscopic and **(E)** microscopic analyzes of the H&E-stained sections of the harvested lungs from dead mice at the oral administrated group showed abscess lesions formation. **(F)** Also, these lungs were fulfilled by macrophages. (*: *P* <0.05, ns: not significant)

Safety is one of the main concerns of utilizing bacteria for cancer treatment due to the colonization of normal tissues and systemic infection^6^. Therefore, the safety of i.v administration of *V. parvula* as the most effective administration route (according to tumor colonization parameter) was evaluated by blood biochemistry and histopathological assays. Healthy BALB/c mice (n=5) were i.v injected with 1 × 10^7^ and sacrificed after 30 days. Macroscopic examination of all the vital organs including liver, kidneys, lungs, and brain did not show any changes as compared to the control mice organs, except the spleen. Apparent splenomegaly was observed in *V. parvula* treated mice (Figure 2A & B). Significant hyperplasia in the splenic white pulp of *V. parvula* treated mice was observed compared with control (Figure 2C & D). The white palp expansion is usually in response to antigenic stimulation of the immune system and causes significant spleen enlargement^38,39^. Furthermore, plasma levels of AST and ALT were measured for liver function analysis. An increase of these enzymes’ plasma levels is considered as a key marker of hepatocellular damage^40^. As Figure 3A illustrates, both of these biomarkers’ levels were in the normal range as compared to the control group. This fact was the same for the creatinine as a well-known kidney function biomarker^41^ (Figure 3A). Besides, no sign of inflammation, abscess lesion formation, parenchymal architecture change, or fibrosis was observed in these organs according to histopathological analyzes (Figure 2B). Taking together, *V. parvula* has a considerable ability for tumor-specific colonization. It can significantly colonize at the tumor after i.v administration, with no significant side effects on the normal organs. Also, *V. parvula* colonization at the tumor site was evaluated weekly for 4 weeks after i.v administration. The bacteria remained detectable (CFUs > 10^6^) at the tumor tissue within these 4 weeks (data not shown).

**Figure 2:**
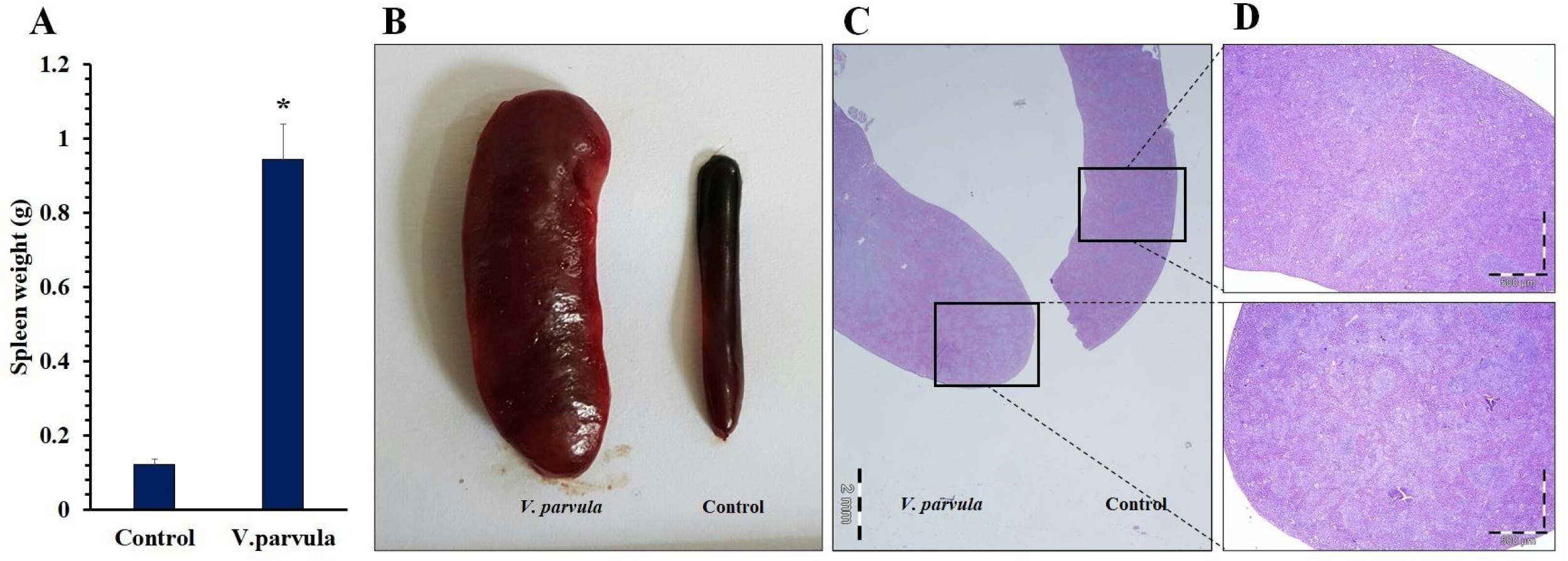
Spleen enlargement (splenomegaly) after i.v administration of *V. parvula* to healthy BALB/c mice. **(A)** Mean spleen weights comparison between the control (injected with PBS) and *V. parvula* (10^7^) i.v administered mice (n=5). **(B)** Macroscopic view of two spleens from the control (right) and *V. parvula* i.v injected mice (Left). H&E-stained sections of spleens from the control (right) and *V. parvula* i.v injected mice (Left) under **(C)** low and (×4) **(D)** high magnifications (×40).

**Figure 3:**
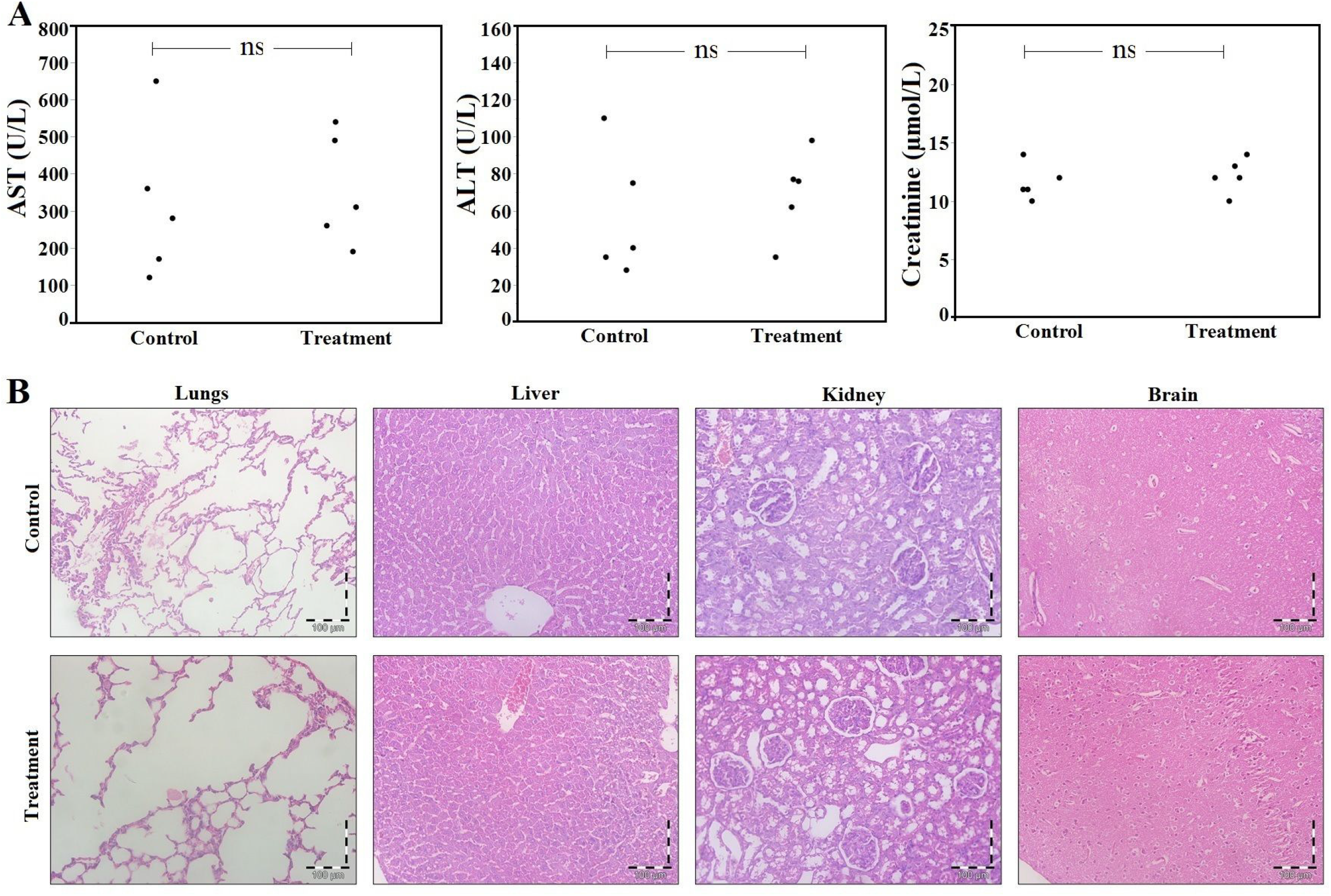
**(A)** Blood biochemical and **(B)** histopathological analyzes of non-tumor bearing mice (n=5), 30 days after i.v administration of 10^7^ *V. parvula* in comparison with control.

### 3.3. Macroscopic and microscopic changes at the breast tumor after *V. parvula* colonization

To investigate the macroscopic and microscopic effects of *V. Parvula* colonization at the breast tumors, the tumor-bearing mice (n=5, mean tumors’ volume: 368 ± 80 mm^3^) were sacrificed 3 days after i.v administration of 10^7^ *V. parvula*. As Figure 4A illustrates, some tumors at the V. parvula administered group exhibited pus formation which contained a high concentration of alive *V. parvula* according to gram staining and culturing the pus samples. Also, tumor H&E-stained sections exhibited clustered colonies of *V. parvula* at the central regions of the tumor (Figure 4B & C) which is mostly hypoxic and necrotic regions. The peripheral regions were free of any colonies of bacteria (Figure 4D). Taking together, multiple clustered colonies of *V. parvula* was observed at the central regions of 4T1 breast tumors which demonstrates high efficacy o this anerobic species to colonize at the hypoxic regions of solid tumors.

**Figure 4:**
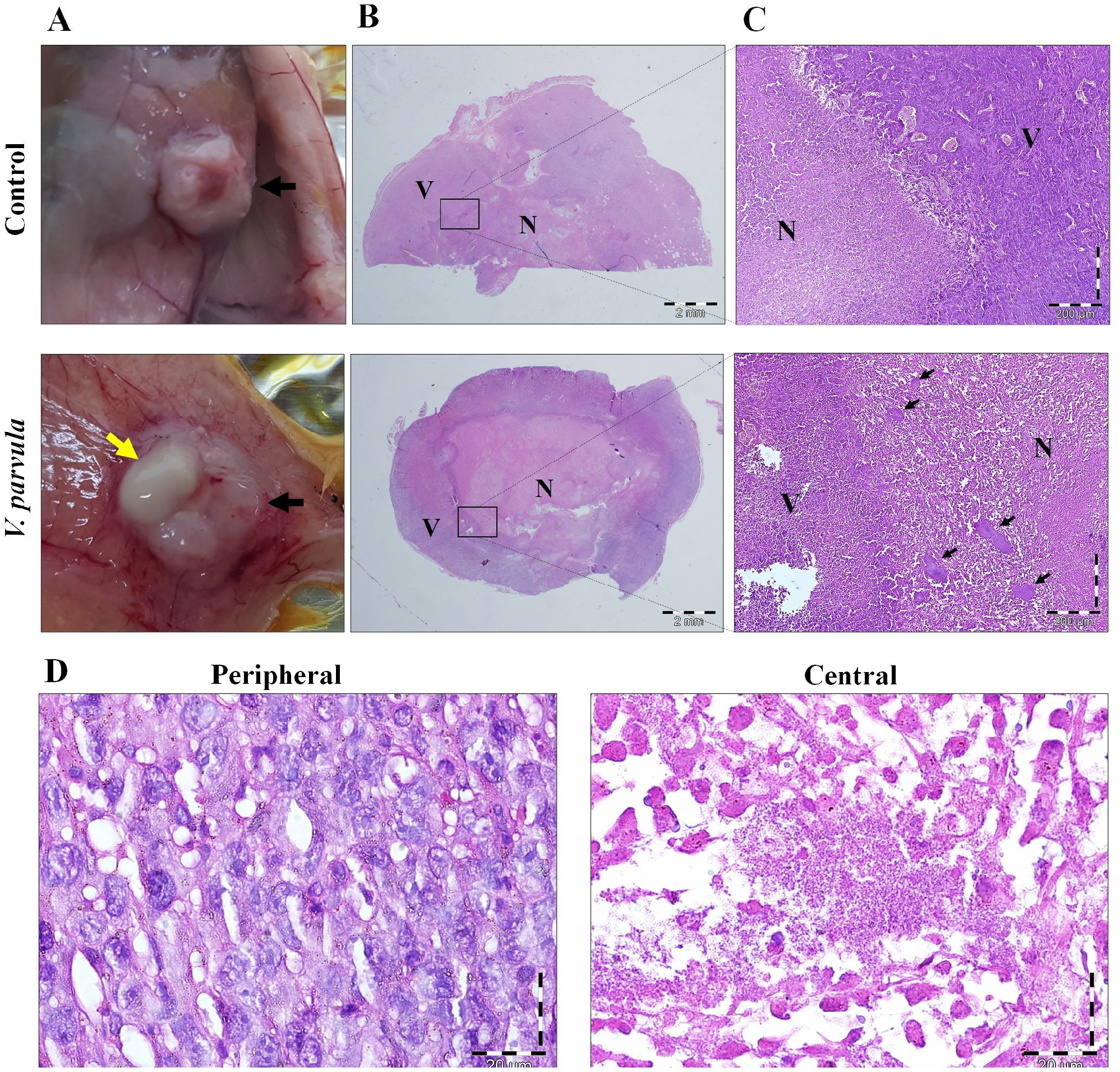
4T1 breast tumor colonization by *V. parvula* after i.v administration. (**A**) Macroscopic view of 4T1 tumors in the control and *V. parvula* administered tumor-bearing mice (72 h after i.v administration of 10^7^ *V. parvula*, n=5, mean tumors’ volume: 368 ± 80 mm^3^). Some of the tumors exhibited pus formation (Yellow arrow: pus, black arrow: tumor). **(B)** Low (×4) and (**C**) high-power (×100) field microscopic view of the H&E-stained sections of 4T1 breast tumors of the control and *V. parvula* administered groups (N: necrosis, V: viable, black arrows indicate the clustered colonies of *V. parvula*). (**D**) High-power (×400) field microscopic view of the central and peripheral regions of the H&E-stained sections of 4T1 breast tumors of *V. parvula* administered groups (N: necrosis, V: viable). The colonies of *V. parvula* were just observed at the central regions of the tumors and peripheral regions were clear from bacteria.

### 3.4. Effect of *V. parvula* colonization on the 4T1 tumors’ growth progression, metastasis, and tumor-bearing mice survival

4T1 breast tumor-bearing mice were i.v injected with 10^7^ *V. parvula* on the 9^th^ day after the cancer cells implantation. The mean tumor volume was 109 ± 18.6 mm^3^. The *V. Parvula* colonization at the tumor site caused significant inhibition (P <0.05) of the tumor growth progression at the first days after administration in comparison with control. However, *V. parvula* colonization couldn’t maintain its therapeutic effect and the mean tumor volume progression reached the control group on the last day of tumors’ growth monitoring. Although a 25.4% decrease in the mean tumors’ volume was observed at the last days of monitoring, the difference between the control and *V. parvula* groups was not statistically significant (P > 0.05) (Figure 5A). Besides, tumor colonization of *V. parvula* didn’t cause an increase in the tumor-bearing mice survival time in comparison with control (Figure 5B). Moreover, no significant therapeutic effect was observed at the breast tumor’s metastasis according to the evaluation of metastatic colonies at the H&E-stained sections of these two groups mice livers (Figure 5C & D). In general, although V. parvula exhibits high tumor targeting and colonization efficacy, its anti-tumor effects is transient.

**Figure 5:**
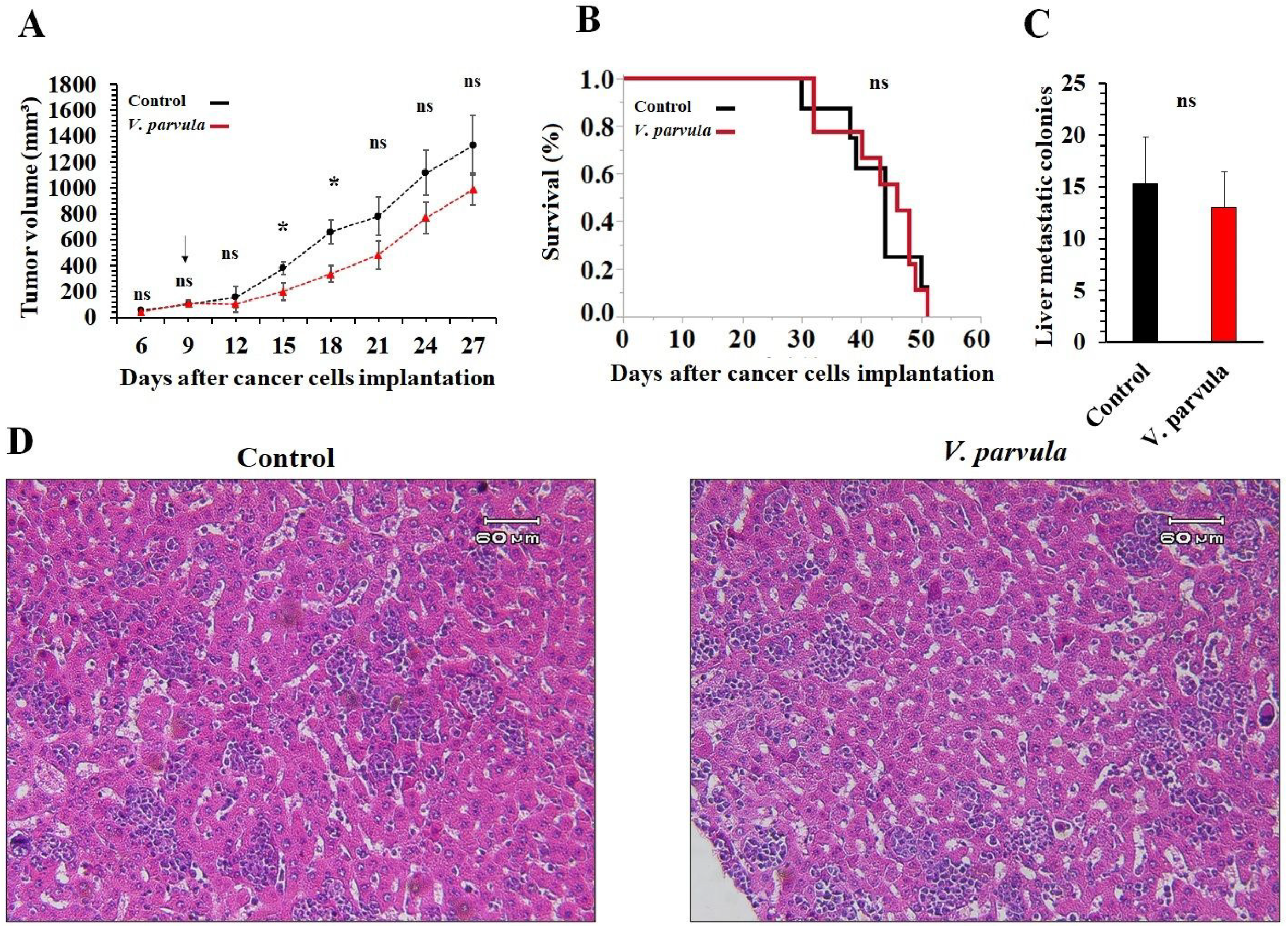
The therapeutic effects of i.v administration of *V. parvula* on the 4T1 breast tumors’ growth progression, metastasis, and tumor-bearing mice survival time. 4T1 tumor-bearing mice were i.v injected with 10^7^ *V. parvula* on the 9^th^ day after cancer cells implantation (Mean tumors volume: 109 ± 18.6 mm^3^). The black arrow indicates the *V. parvula* injection day. (**A**) The mean tumors’ volume progression cures of the control and *V. parvula* groups (*: P < 0.05, ns: not significant). (**B**) Kaplan-Meier survival curves of the control and *V. parvula* groups. (**C**) The average number of metastatic colonies at 10 random high-power (× 200) microscopic fields of the H&E-stained sections of the liver at 40^th^ day after cancer cell implantation (n=5). (**D**) Two sample microscopic field (× 200) of the H&E-stained sections of liver from the control and *V. parvula* administered groups (arrows indicate metastatic colonies).

## Conclusions

Bacteria therapy has gained lots of attention for cancer treatment. Different species and their attenuated strains have entered clinical trials for cancer treatment. In the present study, a novel strictly anaerobic bacterial species was intravenously administrated into 4T1 breast tumor-bearing mice. *V. parvula* exhibited significant safety and tumor-specific colonization. Also, it exhibited high tumor penetration and colonization at the deep regions of the tumor. However, it couldn’t cause a significant therapeutic effect on the breast tumor growth progression or metastasis.

## Abbreviations

V. parvula: Veillonella parvula
i.v: intravenous
i.p: intraperitoneal
i.t: intratumoral
Colony forming units: CFUs
H&E: hematoxylin and eosin

## Contributions

Dr. Kefayat, Dr. Ghahremani, and Dr. Rostami conceived and designed the experiment. The bacteria culture and preparation were performed by Dr. Rostami. All the animal experiments, data analyses, and manuscript writing were performed by Dr. Kefayat, Dr. Ghahremani, and Dr. Rostami.

## Acknowledgment

The authors thank the Arak University of Medical Sciences for their supports.

